# Low-Intensity Focused Ultrasound Enhances Meningeal Lymphatic Drainage for Preventing Cognitive Decline in Alzheimer’s Disease

**DOI:** 10.64898/2026.04.06.716653

**Authors:** Xingjun Xu, Zhou Feng, Tao Jiang, Caixin Zhu, Yu Tang, Yue Shu, Qining Wang, Xiaoli Li, Jingming Hou

**Affiliations:** Department of Rehabilitation, Southwest Hospital, Third Military Medical University (Army Medical University), Chongqing, 400038, China; Department of Advanced Manufacturing and Robotics, College of Engineering, Peking University, Beijing, 100871, China; Pazhou Laboratory (Guangzhou) & School of Automation Science and Engineering, South China University of Technology, Guangzhou, 510641, China

## Abstract

Meningeal lymphatic vessels (mLVs) are vital for brain waste clearance, making them a promising therapeutic target. However, effective modulation strategies for mLVs with translational potential remain underdeveloped. Here, we develop a low-intensity focused ultrasound (LIFU) strategy that precisely targets the vault cranial meninges to non-invasively facilitate mLVs drainage. Using models of Alzheimer’s disease (AD) and aging, we demonstrate that this approach promotes CSF drainage, prevents cognitive decline, and reduces pathological biomarkers. Mechanistically, RNA sequencing combined with calcium imaging in vitro reveals that LIFU activates the Piezo1 ion channel in lymphatic endothelial cells, whereas pharmacological inhibition of Piezo1 abolishes LIFU’s therapeutic effects. Compliant with FDA safety guidelines, this LIFU protocol demonstrates strong clinical translatability. If its efficacy is clinically confirmed, LIFU offers a promising therapy for neurodegenerative diseases triggered by waste accumulation.

## Introduction

Alzheimer’s disease (AD), the most common type of dementia, constitutes 60–80% of neurodegenerative disorder cases and presents an escalating public health burden, particularly in aging populations(1). AD is characterized by the accumulation of β-amyloid (Aβ) plaques and neurofibrillary tangles, which drive neuronal dysfunction and progressive cognitive decline(1, 2). A traditional viewpoint is that the brain is considered immunologically privileged due to the absence of lymphatic system tissues(3, 4). However, the recent finding of meningeal lymphatic vessels (mLVs) in the dura mater has reshaped understanding of waste clearance in the central nervous system (CNS)(5, 6). Dysfunction in this drainage network, which is essential for clearing toxic metabolites, drives both aging and AD progression(7). Given that enhancing mLVs function (e.g., via vascular endothelial growth factor C (VEGF-C) delivery) improves waste clearance and cognition in preclinical models, mLVs represent a promising therapeutic target for AD(7, 8). The development of a non-invasive physical method to modulate mLVs function to improve waste clearance has become an urgent need in the treatment of AD.

The growth and expansion of lymphatic networks depend on mechanical shear forces generated by interstitial fluid and laminar flow(9, 10). Reduced mechanosensitivity in mLVs has been linked to impaired cerebrospinal fluid (CSF) clearance, suggesting that mechanical stimulation could offer a viable therapeutic strategy(11, 12). A major hypothesis is that endothelial cells (LECs) of meningeal lymphatic vessels express mechanically sensitive ion channels (e.g., Piezo1) that sense mechanical stress(11, 12). After activation of these channels, calcium ion influx occurs in the LEC, triggering downstream signaling pathways (such as MAPK and PI3K/Akt), promoting LEC proliferation, migration, and lymphangiogenesis, enhancing the complexity and connectivity of the lymphatic network(11–14).

Low-intensity focused ultrasound stimulation (LIFU) is a non-invasive neuromodulation technique that offers high spatial resolution and deep penetration(15). It modulates neural activity via mechanical effects at safe, low-energy levels, making it a promising therapeutic tool for neurological disorders(16–18). Previous studies have demonstrated that ultrasound stimulation of brain parenchyma can effectively modulate glymphatic function, promoting CSF circulation and facilitating waste clearance, with considerable therapeutic promise for AD and hemorrhagic stroke(19–24). However, therapeutic focused ultrasound stimulation of brain parenchyma presents certain safety considerations that merit further evaluation.(25). MLVs are mechanosensitive and serve as a key pathway for CSF drainage, rendering them potentially amenable to LIFU modulation. Nevertheless, the modulatory effect of LIFU on mLVs remains unexplored.

While several studies suggest that basal lymphatic vessels play a major role in CSF drainage, the contribution of vault cranial mLVs remains significant due to the presence of arachnoid cuff exits (ACE)(26, 27). Meanwhile, targeting these basal lymphatic vessels using ultrasound poses significant technical challenges. In contrast to the relatively uniform cranial vault, the skull base has an irregular anatomical structure that leads to inconsistent and unpredictable propagation of ultrasound waves(25, 28). Moreover, the skull base is adjacent to vital neuroanatomical structures, including the brainstem. Even at low ultrasound intensities, imprecise focusing carries a risk of thermal or mechanical injury to these delicate regions (25, 28). Given the clinical translation challenges posed by the skull base, we established a safe and feasible LIFU strategy that precisely targets the cranial vault meninges to enhance mLV-mediated CSF drainage without stimulating brain parenchyma.

In this study, we demonstrate that LIFU targeting meninges induces structural and functional improvements in mLVs, alleviates pathological features, and enhances cognitive function in AD and aged mouse models. Mechanistically, we identify that LIFU restores meningeal lymphatic structure and circulation by activating piezo type mechanosensitive ion channel component 1 (Piezo1) ion channels in lymphatic endothelial cells (LECs). These findings support the development of LIFU as a promising non-invasive strategy for enhancing mLVs function and mitigating neurodegenerative disease progression.

## Results

### LIFU allows for safe and non-invasive enhancement of mLVs drainage

To ensure the safety of ultrasonic treatment, we targeted LIFU at the confluence of sinuses (CoS) region, where the brain is located more than 1.0 mm from the scalp, minimizing the risk of direct brain stimulation. Mice were anesthetized, and their dorsal scalps were shaved before undergoing LIFU of the CoS region. We designed a collimator to control the effective longitudinal depth of the LIFU stimulation to be no more than 1.0 mm downward from the surface of the skull, so as to enable the ultrasonic waves to specifically stimulate mLVs (Fig. 1 A–C).

**Figure 1.**
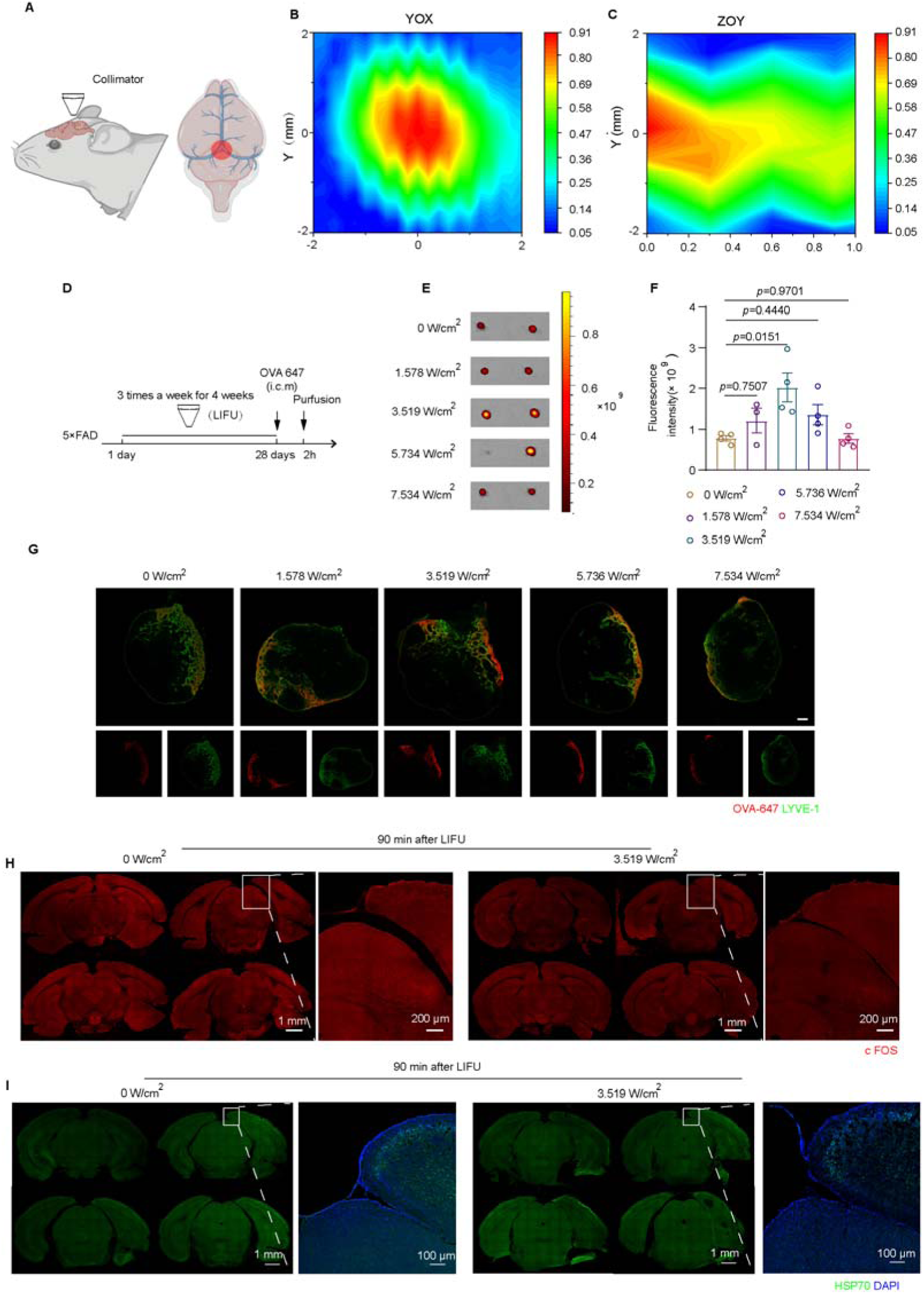
LIFU stimulation protocol for enhancing meningeal lymphatic drainage (5 × FAD, 9-10 month, female) **. A**, Schematic of LIFU targeted stimulation of mLVs. **B**–**C**, Two-dimensional normalized axial (**B**) and lateral (**C**) acoustic intensity distribution of mouse transcranial acoustic field. **D**, Experimental timeline for assessing LIFU-mediated effects on mLVs drainage function. **E**, In vivo fluorescence imaging of dCLNs harvested 2 hours post-injection of OVA-647 (i.c.m) in LIFU-treated mice. Heat maps depict background-subtracted signal intensity. **F**, Quantitative analysis of OVA-647 fluorescence intensity in dCLNs. *n* = 4 mice**. G**, Immunofluorescence staining of LYVE-1 and OVA-647 in dCLN sections at 2 h after injection (i.c.m). Scale bar = 100 μm. **H**, Representative images of coronal sections of brain below CoS region stained with c-Fos. Scale bar = 1 mm. (Bregma −3.4 – −4.4mm). **I**, Representative images of coronal sections of brain below CoS region stained with HSP 70 and DAPI. Scale bar = 1 mm. (Bregma −3.4 – −4.4mm). Values are expressed as mean ± SEM.

To assess the effect of LIFU on mLVs function and determine the optimal ultrasound parameters, we administered varying spatial peak-pulse average intensity (*I_SPPA_*) (0, 0.665, 1.578, 3.519, and 5.736 W/cm^2^) for 4 weeks to 5×FAD mice (9-10month, female). The deep cervical lymph nodes (dCLNs) were harvested for analysis of two hours post OVA-647 cisterna magna injection (i.c.m) (Fig. 1 D). CSF drainage was assessed by measuring the fluorescence intensity of OVA-647 in the dCLNs, with higher fluorescence indicating enhanced CSF clearance. Both IVIS imaging and immunofluorescence results revealed that LIFU-treated mice had significantly greater distribution of OVA-647 in the dCLNs compared to untreated mice, suggesting that LIFU improved mLVs function and facilitated CSF efflux (Fig. 1 E–G). Among the different ultrasound intensities tested, LIFU at 3.519 W/cm^2^ produced the most pronounced enhancement of mLVs drainage (Fig. 1 E–G). Consequently, this intensity was selected for further investigation in subsequent vivo experiments.

To confirm that our LIFU approach did not stimulate the brain, we performed immunofluorescence staining for c-Fos on coronal brain sections below the CoS region. No activation of c-Fos was observed in the cerebral cortex of the LIFU-treated mice compared to the unstimulated group, indicating that LIFU selectively targeted mLVs without affecting the brain parenchyma (Fig. 1 H). Additionally, to mitigate the risk of thermal effects from ultrasound, we set the ultrasound pulse repetition time (PRT) to 0.5 ms and the ultrasound inter-stimulus interval (ISI) to 10 s. This limits energy deposition per pulse while allowing sufficient heat dissipation. HSP 70 staining of brain parenchymal slices beneath the CoS revealed that 20 minutes of LIFU did not induce thermal effects (Fig. 1 I). In summary, the results suggest our LIFU stimulation protocol for mLVs (LIFU) is safe and able to enhance the drainage function of mLVs in the short term.

### LIFU enhances mLVs drainage and alleviates cognitive decline in AD mice

To further evaluate whether LIFU applied to the cranial meninges enhances meningeal lymphatic drainage and ameliorates cognitive deficits in AD mice. we used two AD models: 5×FAD transgenic mice (9–10 months, female) and intrahippocampal injection of Aβ1-42 oligomers (2 months, male) (Fig. 2 A). Both IVIS imaging and immunofluorescence analysis of dCLNs showed that the influx of OVA-647 into dCLNs was significantly increased in LIFU-treated 5×FAD mice compared with untreated 5×FAD mice (Fig. 2 B–E). To investigate the potential correlation between enhanced lymphatic drainage and mLVs structural remodeling, we performed meningeal lymphatic vessel visualization using LYVE-1 immunofluorescence staining. The results suggest that four weeks of LIFU can enhance the coverage, diameter, and sprout count of LYVE-1^+^ vessels in 5×FAD mice (Fig. 2 F and G).

**Figure 2.**
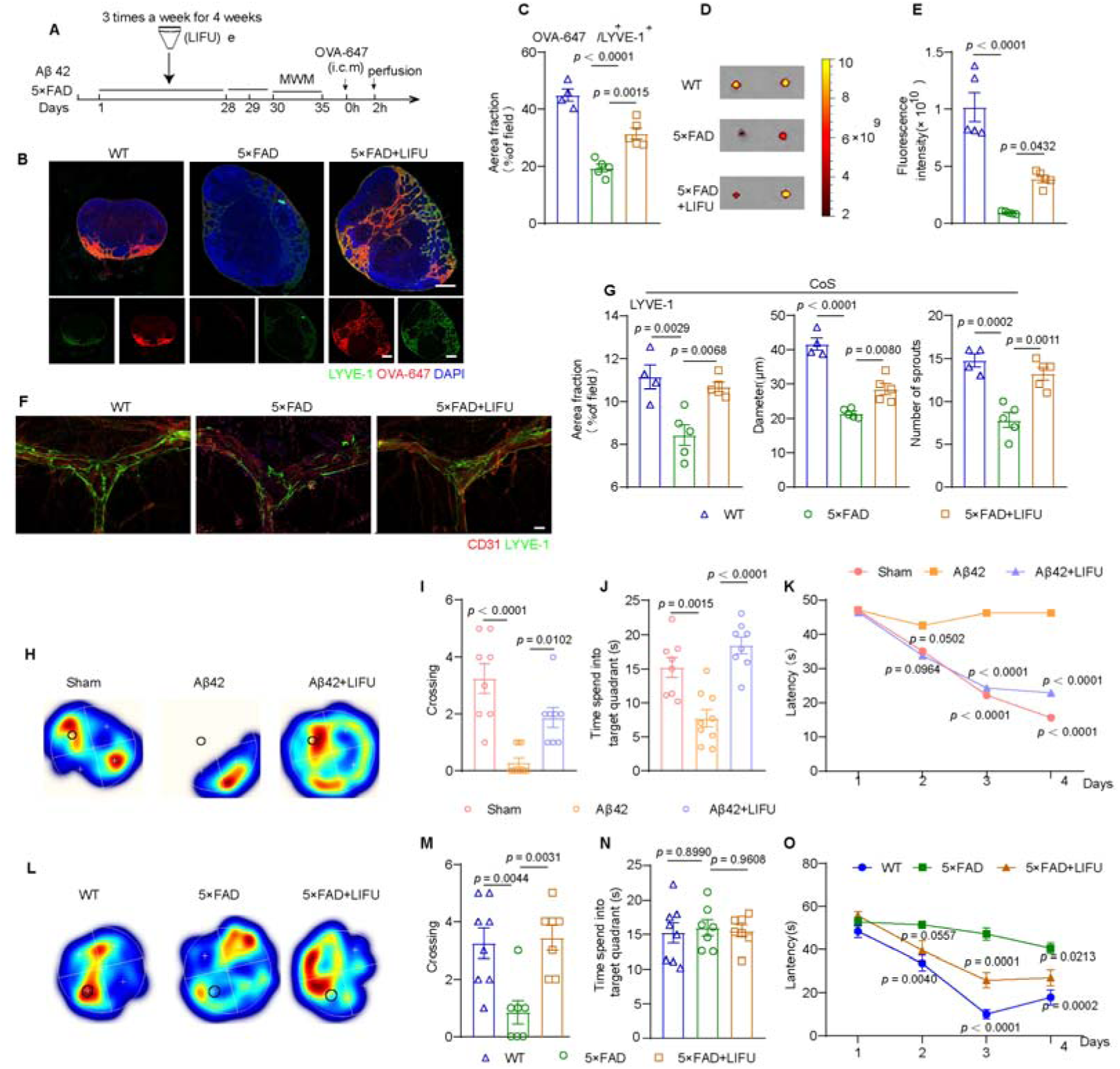
Effects of LIFU on mLVs drainage and cognition in AD mice. **A**, Treatment and behavior test schedule for AD mice. **B**, Representative images of dCLNs from 5×FAD mice (9–10 months) stained with LYVE-1 and DAPI, showing OVA-647 accumulation 2 hours post injection (i.c.m). Scale bar = 200 μm. **C**, OVA-647 fluorescence distribution in dCLN sections in 5×FAD mice. *n =* 4–5. **D**, dCLNs collected 2 hours after OVA-647 injection (i.c.m) in 5×FAD mice, imaged via IVIS epifluorescence. Representative background-subtracted heatmaps. **E**, dCLN fluorescence intensity in 5×FAD mice. *n =* 5 mice**. F**, Representative meningeal images stained with LYVE-1 and CD31 in ×FAD mice (9–10 months). Scale bar = 400 μm. **G**, LYVE-1^+^ lymphatic vessel area fraction, sprout count, and mLVs diameter in the confluence of sinuses (CoS) region. *n =* 4–5 mice. **H**–**K**, Representative occupancy heatmaps (**H**), platform crossing times (**I**), time spent in target quadrant (**J**), and latency to reach the platform (**K**) from MWM tests in Aβ42-injected mice. *n =*8–12 mice. **L**–**O**, Occupancy heatmaps (**L**), platform crossings (**M**), target quadrant time (**N**), and latency from MWM tests (**O**) in 5×FAD mice (9–10 months). *n =*7–8 mice. Aβ42-injected mice: Aβ42. Data in **C**, **E**, **G, I**–**K** and **M**–**N** are shown as mean ± SEM.

We then assessed the cognitive effects of mLVs modulation via LIFU in AD mice. After 4 weeks of LIFU treatment, cognitive function was evaluated the Morris’s water maze (MWM) tests (Fig. 2A). In the MWM test, both non-LIFU 5×FAD and non-LIFU Aβ42-injected mice exhibited longer escape latencies and fewer platform crossings, indicative of impaired long-term spatial memory. LIFU treatment significantly improved performance in these measures, recovering spatial memory deficits in AD mice (Fig. 2 H-O). These findings collectively demonstrate LIFU alleviates cognitive impairments, particularly in spatial learning and memory, in AD mice.

### LIFU attenuates pathological damage in AD mice

Aβ accumulation, neuroinflammation, and neuronal damage are key pathological features of AD(1). To assess these features in 5×FAD mice, we performed immunostaining for Aβ (D54D2) and Iba1 (a microglial marker). Results showed significant amyloid plaque deposition throughout the brain of 5×FAD mice, which was reduced in the hippocampus (HPC) and prefrontal cortex (PFC) of LIFU-treated 5×FAD mice (Fig. 3 A–D, I–K). Additionally, analysis of Iba1^+^ cell coverage and the number of peri-Aβ plaque Iba1^+^ cells in the HPC revealed that LIFU treatment alleviated neuroinflammation in 5×FAD mice (Fig. 3 A, F, and G). To further investigate the neuroprotective effects of LIFU, we performed immunostaining with NeuN, a neuronal marker in HPC. Compared to WT littermates, non-LIFU 5×FAD mice showed a significant reduction in NeuN^+^ neurons, which was partially restored by LIFU treatment, bringing neuron numbers closer to those observed in WT controls (Fig. 3 E and H). These findings indicate that LIFU mitigates AD pathology by reducing Aβ accumulation, suppressing neuroinflammation, and promoting neuronal preservation, ultimately fostering a remediated cerebral microenvironment.

**Figure 3.**
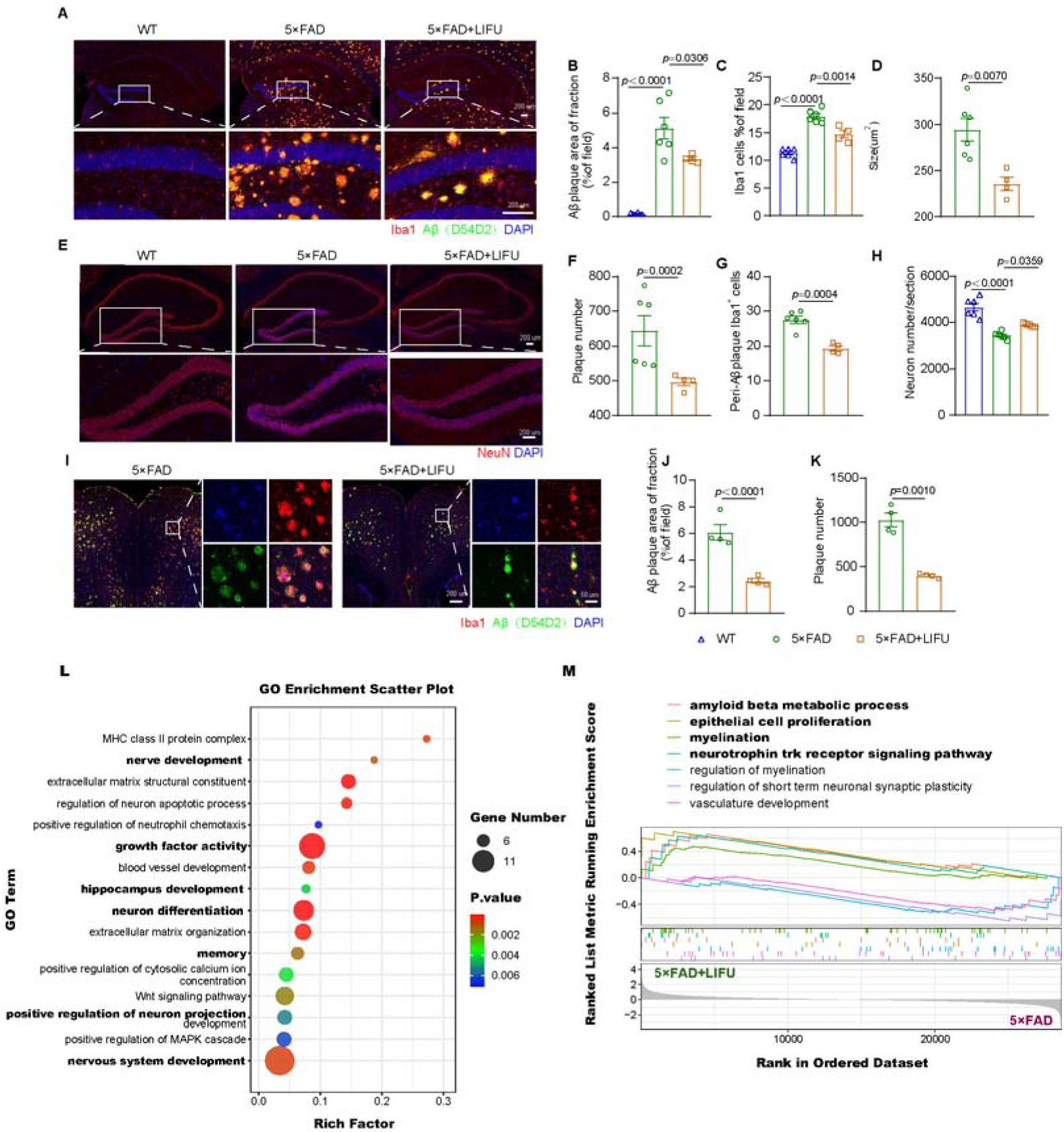
Effects of LIFU on pathology in 5×FAD (9–10month) mice. **A**, Representative images of HPC brain sections stained with Aβ(D54D2), Iba1, and DAPI (2–3 replicates). Scale bar =200 μm. **B**–**G**, Aβ plaque area fraction (**B**), plaque size (**C**), and plaque number (**D**), Iba1^+^ area fraction (**F**), and Iba1^+^ cells per Aβ plaque (**G**) in HPC. *n =*4–6 mice. **E**, Representative HPC sections stained with NeuN and DAPI (2–3 replicates). Scale bar = 200 μm. **H**, Neuron count in HPC. *n =*5 mice in each group. **I**, Representative PFC sections stained with Aβ(D54D2), Iba1, and DAPI (2–3 replicates). Scale bar =200 μm or 50 μm. **J**–**K**, Aβ plaque area fraction (**J**) and number (**K**) in PFC. *n =* 4 mice in each group. **L**, GO terms following LIFU treatment in HPC of 5×FAD mice. *n =* 3 mice in each group. **M**, GSEA enrichment plots. *n* = 3 mice in each group. Data in **B**–**D**, **F**–**H**, and **J**–**K** are shown as mean ± SEM. Two-tailed t-tests analyzed data in **C**, **D**, **G**, **J**, and **K**, while one-way ANOVA with Sidak’s multiple comparison test examined data in **B**, **F**, and **H** for multi-group comparisons. In **L**, *P* values used the hypergeometric test via clusterProfiler, and in **M**, an empirical phenotype-based permutation test calculated *P* values; neither adjusted for multiple comparisons.

To further evaluate LIFU’s impact on AD-related pathological changes, we conducted hippocampal transcriptomic profiling in 5×FAD mice. Total RNA was extracted from hippocampal tissue and sequenced. Gene Ontology (GO) functional enrichment analysis revealed significant changes in gene sets related to nerve development, neuron projection, memory, and nervous system development (Fig. 3 L). Hallmark gene set-based gene set enrichment analysis (GSEA) revealed significant upregulation of biological pathways relating to AD pathology amelioration, including amyloid-β metabolism, epithelial cell proliferation, myelination, and neurotrophin receptor signaling. (Fig. 3 M). These results suggest that LIFU treatment may modulate amyloid beta metabolism, neuronal survival, and other AD-related pathways, contributing to the amelioration of AD pathology.

### LIFU enhances mLVs drainage and alleviates cognitive decline in aged mice

To further validate our findings in AD mice, we conducted a similar study in aged mice (18–19 months, male) and assessed the effects of LIFU treatment on cognitive function (Fig. 4 A). The WMZ test was used to evaluate changes in cognitive function after 4 weeks of LIFU treatment. Long-term spatial memory was assessed using the MWM test. Non-LIFU aged mice exhibited significant impairments in long-term contextual memory compared to the younger cohort, while LIFU treatment improved performance (Fig. 4 B–E). In conclusion, these behavioral test results demonstrate that 4 weeks of LIFU treatment alleviates anxiety-like behaviors and improves cognitive function in 18–19-month-old mice.

**Figure 4.**
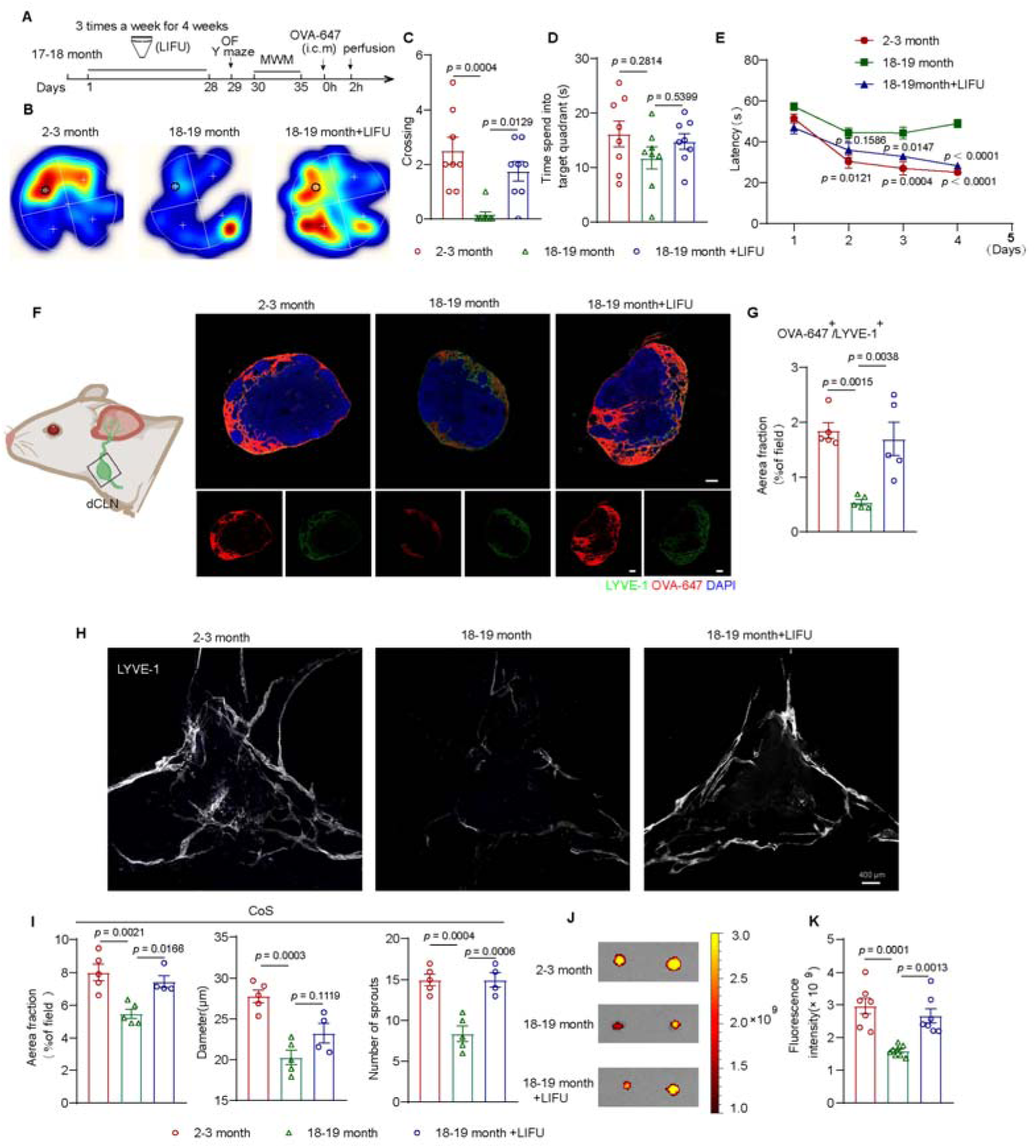
Effects of LIFU on cognition and mLVs drainage in aged (18–19 month) mice. **A**, Treatment and behavior test schedule for aged mice. **B**–**E**, Representative occupancy heatmaps **(B)**, platform crossing times **(C)**, time spent in target quadrant **(D)**, and latency to reach platform **(E)** from the MWM test. *n* =8 mice in each group. **F**, Representative dCLN sections stained with LYVE-1 and DAPI, showing OVA-647 accumulation 2 hours post injection (i.c.m). Scale bar = 200 μm. **G**, OVA-647 fluorescence distribution in dCLN sections. *n =* 5 mice in each group. **H**, Representative images of meninges stained with LYVE-1. Scale bar = 400 μm. **I**, LYVE-1^+^ lymphatic vessel area fraction, sprout count, and mLVs diameter in CoS. *n =* 4–5 mice. **J**, dCLNs collected 2 hours after OVA-647 injection (i.c.m), imaged via IVIS epifluorescence. Representative background-subtracted heatmaps. **K**, dCLNs fluorescence intensity. *n =* 7–8 mice. Data in **C**–**E**, **G**, **I**, and **K** are shown as mean ± SEM.

We then explored whether 4 weeks of LIFU treatment could also regulate the function of the meningeal lymphatic system in 18–19-month-old mice. Two hours after the injection (i.c.m.) of OVA-647, we harvested the dCLNs of the mice and performed IVIS imaging and immunofluorescence staining. The results from the dCLNs showed that, compared with 2–3-month-old mice, non-LIFU 18–19-month-old mice had drainage disorders in the meningeal lymphatic system. However, LIFU treatment alleviated the drainage disorders of the lymphatic system in 18–19-month-old mice (Fig. 4 F, G, J, and K).

To detect whether there were structural changes in the meningeal lymphatic vessels after LIFU treatment, we performed immunofluorescence staining of LYVE-1 on the meninges and statistically analyzed the LYVE-1 at the CoS. The results showed that, compared with 2–3-month-old mice, non-LIFU 18–19-month-old mice exhibited abnormal mLVs structures. Specifically, the area fraction of LYVE-1^+^ at the CoS, the diameter of the meningeal lymphatic vessels, and the number of sprouts were lower. LIFU reduced these structural abnormalities at the CoS in 18–19-month-old mice (Fig. 4 H and I). The above results suggest that LIFU improves lymphatic drainage function and reduces structural abnormalities in the meningeal lymphatic system in aged mice, supporting the results observed in AD mice.

### Enhanced mLVs drainage is essential for cognitive improvement induced by LIFU

To investigate whether LIFU’s cognitive benefits and amelioration of AD pathology depended on improved mLVs drainage, we performed mLVs ablation in three mouse models: aged (18-19 months, male), Aβ42-injected, and 5×FAD (9-10 months, female) mice, following established protocols (Figs. 5A, 6A). LIFU administration enhanced learning and memory performance across all three models, demonstrating its therapeutic potential for age-related and AD-associated cognitive decline highlighting the critical role of mLVs in cognitive recovery via LIFU treatment (Fig. 5 B–I, Fig. 6 B–H).

**Figure 5.**
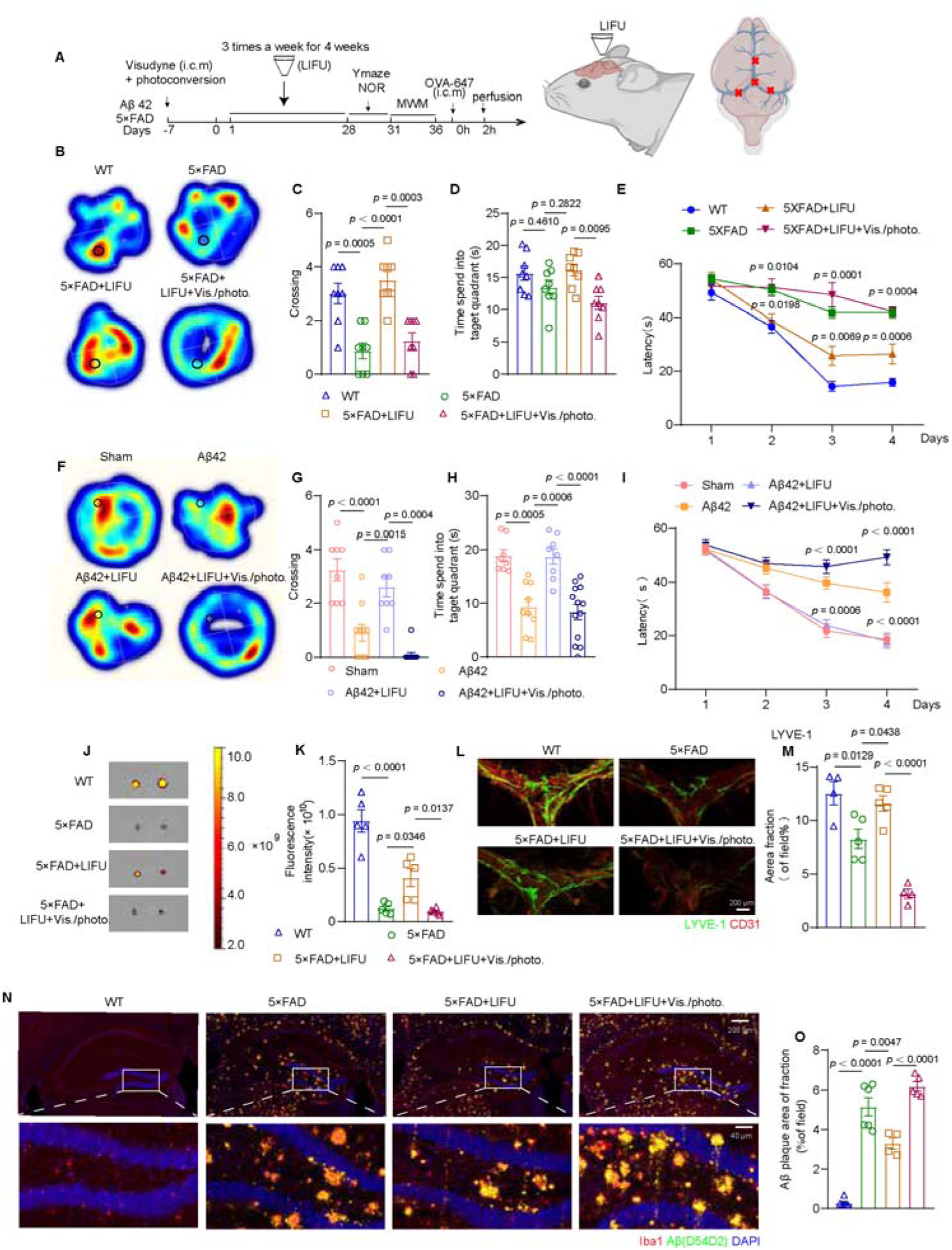
Effects of LIFU on cognition and pathology in mLVs-ablated AD mice. **A**, Treatment and behavior test schedule for mLVs-ablated AD mice. **B**–**E**, Representative occupancy heatmaps (**B**), platform crossing times (**C**), time spent in target quadrant (**D**), and latency to platform (**E**) from MWM tests in mLVs-ablated 5×FAD (9–10 month) mice. *n =* 7–8 mice. **F-I**, representative occupancy heatmaps **(F)**, platform crossing times **(G)**, time spent in target quadrant **(H)**, and latency to platform **(I)** from MWM tests in mLVs-ablated Aβ42-injected mice. *n =*8–12 mice. **J**, dCLNs collected 2h after OVA-647 injection (i.c.m) in mLVs-ablated 5×FAD mice, imaged via IVIS epifluorescence. Representative background-subtracted heatmaps. **K**, dCLNs fluorescence intensity. *n =* 7–8 mice**. L**, Representative meningeal images stained with LYVE-1 and CD31 in mLVs-ablated 5×FAD mice. Scale bar = 100 μm. **M**, LYVE-1^+^ lymphatic vessel area fraction from CoS. *n =* 4–5 mice. **N**, Representative HPC brain sections stained with Aβ(D54D2), Iba1, and DAPI (from 2–3 replicates) in mLVs-ablated 5×FAD mice. Scale bar = 200 μm or 40 μm. **O**, Aβ(D54D2) area fraction in HPC in mLVs-ablated 5×FAD mice. *n =* 4–6 mice. Aβ42-injected mice: Aβ42. Data in **C**–**E**, **G**–**I**, **K**, **M**, and **O** are shown as mean ± SEM.

**Figure 6.**
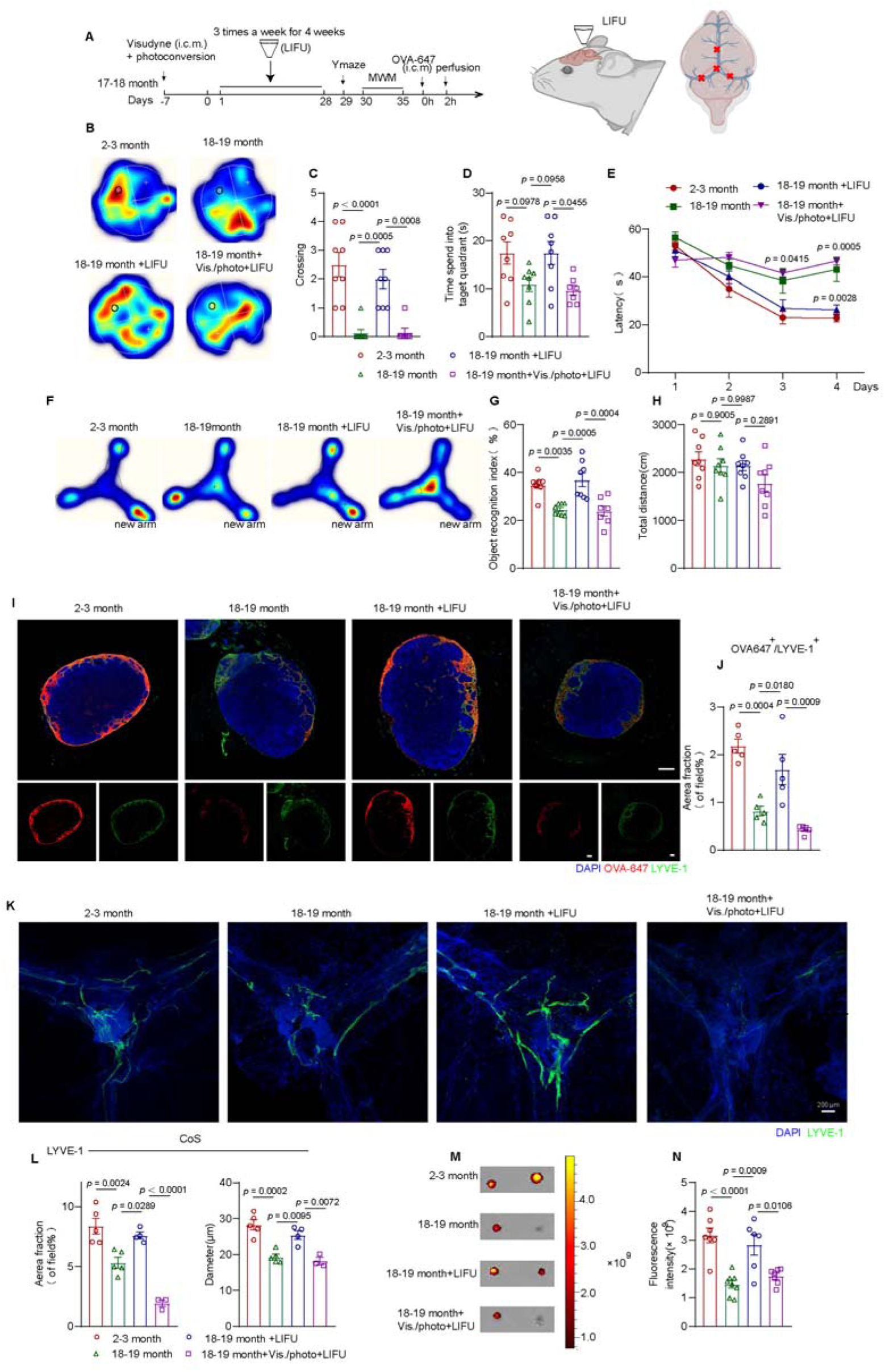
Effects of LIFU on cognition and lymphatic drainage in mLVs-ablated aged (18–19 month) mice. **A**, Treatment and behavior test schedule for mLVs-ablated aged mice. **B**–**E**, Representative occupancy heatmaps (**B**), platform crossing times (**C**), time in target quadrant (**D**), and latency to platform (**E**) from MWM tests of mLVs-ablated aged mice. *n =* 8 mice in each group. **F**–**H**, Representative occupancy heatmaps (**F**), novel arm preference (**G**), and total distance (**H**) from Y maze tests of mLVs-ablated aged mice. *n =* 8 mice in each group. **I**, Representative dCLN sections stained with LYVE-1 and DAPI showing OVA-647 accumulation 2 h post injection (i.c.m) in mLVs-ablated aged mice. Scale bar = 200 μm. **J**, OVA-647 fluorescence in dCLN sections in mLVs-ablated aged mice. *n =* 5 mice in each group. **K**, Representative meningeal images stained with LYVE-1 and DAPI in mLVs-ablated aged mice. Scale bar = 200 μm. **L**, LYVE-1+ vessel area fraction and mLVs diameter in CoS region in mLVs-ablated aged mice. *n =* 3–5 mice. **M**, dCLNs imaged by IVIS epifluorescence 2 h after OVA-647 injection (i.c.m) in mLVs-ablated aged mice. Representative background-subtracted heatmaps. **N**, dCLN fluorescence intensity in mLVs-ablated aged mice. *n =* 7–8 mice. Data in **C**–**E**, **G**–**H**, **J**, **L**, and **N** are presented as mean ± SEM.

To further determine whether the abolishment of LIFU treatment effects in aged and AD mice with ablated mLVs resulted from impaired mLVs function, we performed IVIS imaging and immunofluorescence staining of dCLNs and mLVs morphology. The results revealed that meningeal lymphatic ablation negated the benefits of LIFU on mLVs drainage function (Fig. 5 J, K, Fig.6 I, J, M, N). Aged mice and 5×FAD mice treated with LIFU stimulation and ablation exhibited a significant reduction in LYVE-1^+^ coverage area in mLVs. (Fig. 5 L, M, Fig. 6 K, L). Similarly, the beneficial effects of LIFU treatment on amyloid load reduction in the hippocampus were also abolished in mLVs-ablated AD mice (Fig. 5 N, O). These results indicated that enhancing mLVs drainage by LIFU is essential for cognition improvement in aged and AD mice.

### LIFU modulates the gating of Piezo1 ion channels on mLECs for enhancing mLVs drainage

LIFU treatment significantly promoted lymphangiogenesis and led to sustained enhancement of meningeal lymphatic drainage function. To investigate the underlying mechanism, we applied 20 minutes of LIFU stimulation to human lymphatic endothelial cells (hLECs) and collected cells within 10 minutes post-stimulation for RNA-seq analysis of differentially expressed genes (DEGs). GO functional enrichment analysis revealed that LIFU stimulation rapidly enriched gene sets associated with cellular calcium ion influx, including calcium ion binding, voltage-gated calcium channel activity, and calcium ion import (Fig. 7 A). Additionally, calcium-related genes such as *CALM1*, *CALM2*, *CALM3*, and *CAMK2D* were significantly upregulated (Fig. 7 B). Among calcium channels, Piezo1, the most mechanosensitive calcium ion channel, was prominently expressed in LECs(29, 30), as confirmed by the Human Protein Atlas single-cell RNA database (as of October 2025), piezo1 expression in LECs was found to be higher than in approximately 90 other cell types. Previous studies have shown that Piezo1 channels are abundant in mLECs, where they sense CSF**-**derived shear stress and contribute to mLVs development and maintenance(11, 12). Recent evidence indicates that LIFU can activate Piezo1 channels in multiple cell types, including neurons, fibroblasts, and other mechanosensitive cells(31–33). Heatmap analysis further demonstrated that LIFU upregulated genes associated with Piezo1 downstream signaling pathways, including *AKT1*, *ANGPT1,* and *ANGPT2* (Fig. 7 B). Notably, *AKT1*, *ANGPT1*, *ANGPT2*, and *DLL4* are all associated with the proliferation of LECs, and our results revealed that LIFU increased the area and sprout count of mLVs(34–36). Therefore, LIFU may promote lymphatic endothelial cell proliferation through activation of Piezo1 ion channels.

**Figure 7.**
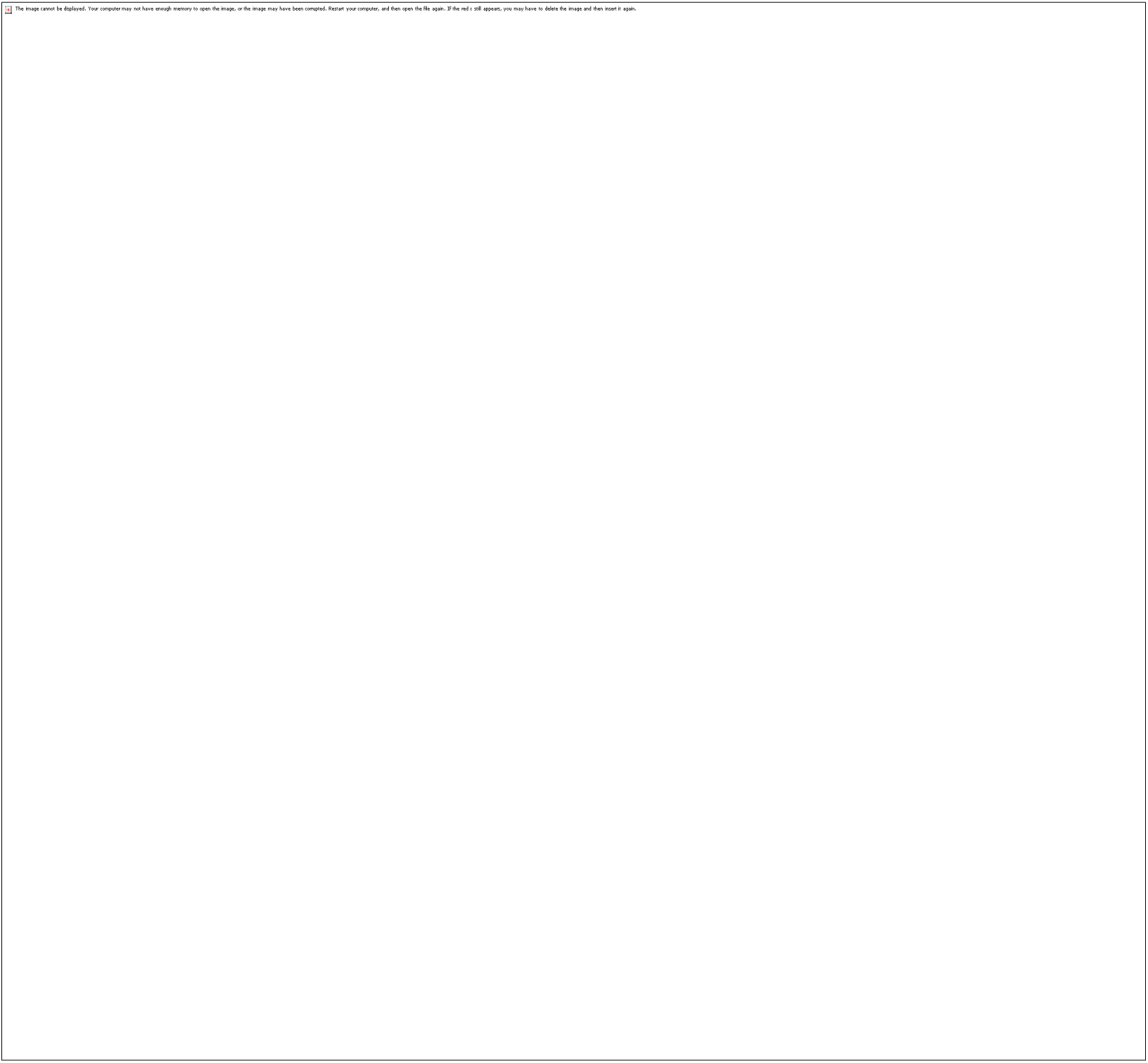
Piezo1 response of mLECs to LIFU stimulation. **A**, GO Enrichment BarPlot upon LIFU in hLECs. **B**, Heatmap showing relative expression levels of DEGs in hLECs involved in calcium ion and cell proliferation. *n* = 3 in each group. **C**, Live cell calcium imaging device. **D**, Fluo-4 AM fluorescence imaging of hLECs on LIFU stimulation. Fluorescence intensity of Fluo-4 reflects concentration of intracellular calcium. Scar bar = 100 μm. **E**, Comparison of fluorescence intensity between baseline and LIFU stimulation periods. *n* = 25 cells in each group. **F**, Representative calcium traces of one hLEC of siNT, GsMTx4, Saline, and siP group. LIFU parameters of hLECs were as follows: frequency: 0.5 MHz, *I_SPPA_*: 1.578 W/cm^2^; stimulation duration: 2 min. siNT: no target siRNA, siP: siRNA-Piezo1. Data in **E** is shown as mean ± SEM.

To validate the results of our RNA-seq analysis, we implemented a comprehensive experimental approach in vitro and in vivo, with genetic and pharmacological interventions, targeting Piezo1 activity. In vitro, we performed live-cell calcium imaging using Fluo-4 AM as an intracellular calcium indicator. The results demonstrated that LIFU stimulation (1.578 W/cm^2^, 0.5 MHz) induced robust calcium transients in control hLECs, which returned to baseline within 2 minutes post-stimulation (Fig. 7 D, E, F). This response was markedly attenuated by either GsMTx4 (a Piezo1 inhibitor) treatment or Piezo1 siRNA knockdown (Fig. 7 D, E, F), confirming the essential role of Piezo1 in mediating LIFU-induced calcium influx in cultured lymphatic endothelial cells.

Piezo 1 is critical for the development and maintenance of the meningeal lymphatic system. Previous studies have shown that activating the Piezo1 ion channel with Yoda 1 promotes CSF drainage, thereby improving diseases characterized by excessive CSF accumulation and mLVs drainage dysfunction, such as hydrocephalus and craniosynostosis(11, 12). Our results demonstrated that LIFU improved mLVs drainage function in AD mice, alleviated AD pathology, and activated Piezo1 ion channels on mLECs. To confirm the role of Piezo1 in LIFU-mediated regulation of the meningeal lymphatic system, we used intrathecal injection of GsMTx4 to inhibit the opening of Piezo1 (Fig. 8 A). Compared to sham-treated 5×FAD mice, LIFU treatment significantly enhanced meningeal lymphatic drainage, increased the LYVE-1^+^area in the CoS, enlarged the diameter of mLVs, and promoted mLVs sprouting. However, these beneficial effects of LIFU on mLVs function were completely abolished following Piezo1 channel inhibition (Fig. 8 B–G). Furthermore, LIFU-induced improvements in microglial activation, amyloid load, and neuroprotection in the HPC was also abolished in GsMTx4-treated 5×FAD mice (Fig. 8 H–N). These results suggest that activation of Piezo1 ion channels is crucial for LIFU-mediated enhancement of mLVs function and its associated therapeutic effects in AD mice.

**Figure 8.**
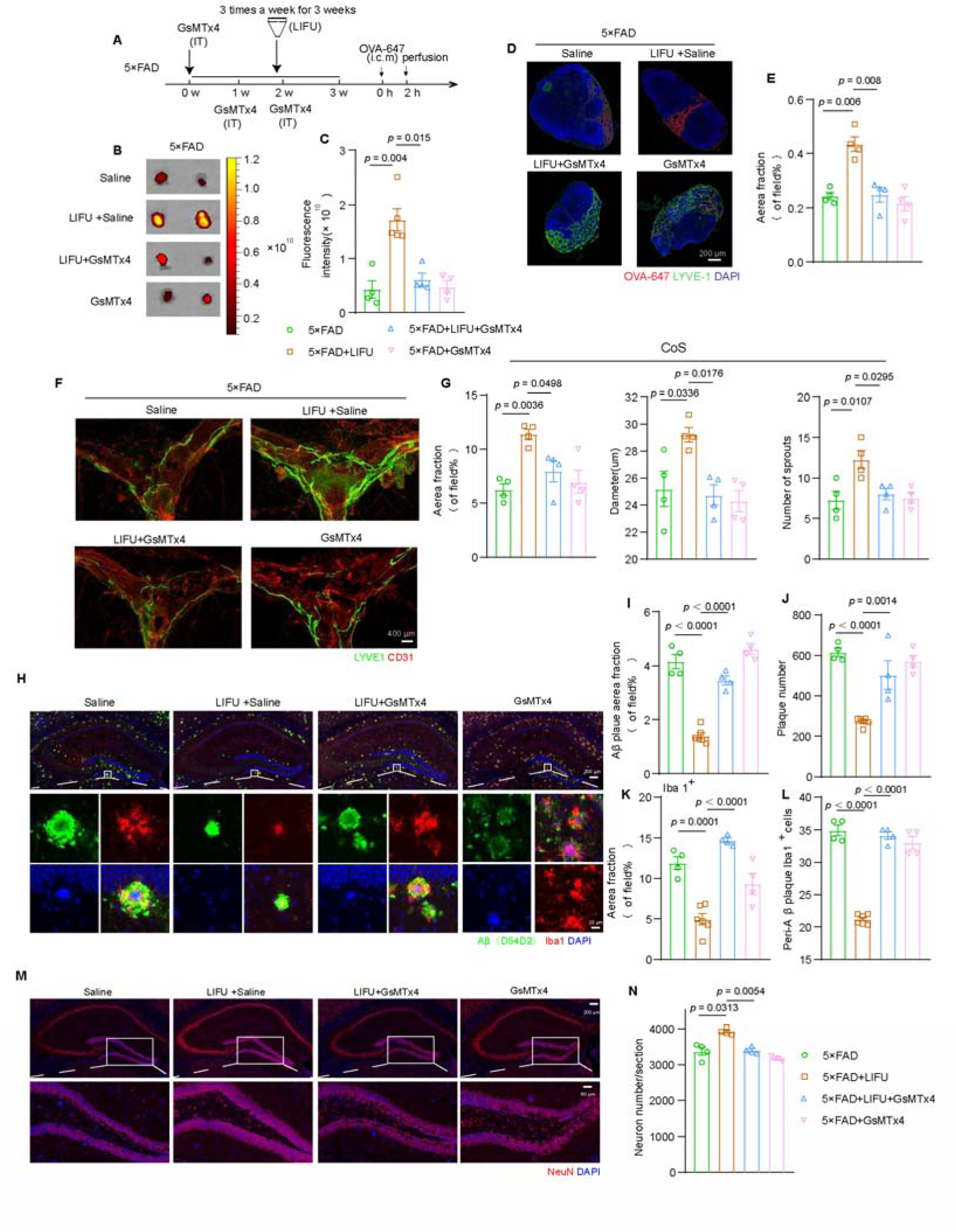
Role of Piezo1 in regulation of mLVs drainage function by LIFU. **A**, Schedule of treatments and GsMTx4 injection (IT) of 5×FAD (9–10 month) mice. **B**, dCLNs collected 2h after OVA-647 injection (i.c.m) in 5×FAD (9–10 month) mice, imaged via IVIS epifluorescence. Representative background-subtracted heatmaps. **C**, dCLNs fluorescence intensity. *n =* 4-5 mice in each group. **D**, Representative section images of dCLNs from 5×FAD mice, which were stained with LYVE-1 and DAPI at 2 hours after OVA-647 injection (i.c.m) to show OVA-647 accumulation. Scale bar = 200 μm. **E**, Fluorescence distribution of OVA-647 in dCLN sections. *n =* 4 mice in each group. **F**, Representative images of the meninges from 5×FAD mice, stained with LYVE-1 and CD31. Scale bar = 400 μm. **G**, Area fraction of LYVE-1^+^ lymphatic vessels and diameter of mLVs in CoS. *n =* 4 mice in each group. **H**, Representative images of HPC brain sections from 5×FAD mice, stained with Aβ (D54D2), Iba1, and DAPI (2–3 replicates). Scale bar = 200 μm or 20 μm. **I**–**L**, HPC area fraction of Aβ (D54D2) (**I**), number of Aβ plaques (**J**), area fraction of Iba1^+^ (**K**), and number of Iba1^+^ cells in each Aβ plaque (**L**). *n =*4 mice in each group. **M**, Representative images of HPC brain sections from 5×FAD mice, stained with NeuN and DAPI (2–3 replicates). Scale bar = 200 μm or 60 μm. **N**, Neuron count in HPC. *n =*4 mice in each group. Data in **C**, **E**, **G**, **I**, **J**, **K**, **L**, and **N** are shown as mean ± SEM.

## Discussion

In this study, we developed a LIFU strategy that precisely targets the vault cranial meninges to non-invasively facilitate mLVs drainage, thereby alleviating cognitive impairments in two AD models (5×FAD and Aβ42-injected mice) and aged mice by enhancing mLVs drainage function. We further demonstrated that LIFU recovered mLVs drainage function and structural damage in AD mice by modulating Piezo1 on mLECs, leading to increased lymphatic clearance of Aβ deposition and reduced neuroinflammation, thereby reducing AD pathology and improving cognition.

By using a collimator to restrict the effective depth of LIFU stimulation to within 1.0 mm, we were able to target the cranial meninges in the CoS while avoiding stimulation of brain parenchyma. Our results demonstrate that 4 weeks LIFU at 3.519W/cm^2^ (*I_SPPA_*) intensity exhibits optimal enhancement of mLVs drainage function in 5×FAD mice. Immunofluorescence analysis of c-Fos in brain coronal sections showed no significant activation of c-Fos in the cortical parenchyma after LIFU stimulation. Given bone tissue’s high ultrasound absorption coefficient, we employed LIFU with 0.5-ms PRT and 10-second ISI. This approach significantly limits per-pulse energy deposition while enabling adequate heat dissipation. Immunofluorescence analysis of HSP70 expression in brain parenchyma beneath the CoS at 90 minutes post-LIFU stimulation confirmed the absence of detectable thermal effects in neural tissue. Laser speckle imaging further revealed that the same 4-week LIFU treatment did not induce alterations in cerebral blood flow in 5×FAD mice. These findings suggest that LIFU, when targeted at mLVs, is a safe and effective method for modulating meningeal lymphatic drainage, offering a promising strategy to enhance lymphatic function without direct stimulation of the brain parenchyma.

Our findings demonstrate that LIFU not only ameliorated drainage dysfunction in both AD and aged mice but also significantly increased the distribution area, sprouting, and structural remodeling of mLVs, culminating in improved cognitive function. Critically, this study establishes that LIFU reduces amyloid plaque burden and neuronal loss in two distinct AD mouse models: hippocampal Aβ42 oligomer-injected mice and 5×FAD transgenic mice. Supporting these observations, RNA sequencing analysis of hippocampi from LIFU-treated 5×FAD mice revealed upregulation of pathways associated with amyloid clearance and neuroprotection. This aligns with previous studies demonstrating that augmenting mLVs drainage through methods such as intracerebral VEGF-C injection or near-infrared light stimulation alleviates cognitive impairment and AD pathology(7, 37, 38). Importantly, our findings confirm that LIFU treatment restores mLVs circulatory function, mitigates AD pathology, and ameliorates cognitive deficits. Consequently, this approach holds significant translational potential for clinical treatment neurological disorders, including AD, in which meningeal lymphatic dysfunction is implicated.

Mechanistically, photodisruption of dorsal mLVs abolished the therapeutic benefits of LIFU in both AD and aged mice, thereby establishing that LIFU efficacy is dependent on enhanced mLVs function. Although cranial base/nasopharyngeal lymphatic plexus regions dominate lymphatic drainage, the discovery of ACE points (peri-bridging venous spaces with leaky barriers) highlights the functional significance of dorsal mLVs in CSF clearance(39–41). Given anatomical context, we focused LIFU on CoS as a single site for practical and safety. Nevertheless, multi-target dorsal mLVs stimulation or skull-base targeting merits exploration to further increase drainage efficacy. For clinical translation, two key challenges require attention: 1) significant ultrasound attenuation through thick skull bone necessitates rigorous parameter optimization; 2) therapeutic synergy should be investigated. Specifically, LIFU represents a promising strategy for AD and related disorders via precise mLVs modulation, and its clinical implementation could synergize with established AD therapies like anti-Aβ immunotherapy to potentially enhance treatment efficacy(8).

Although ultrasound has been shown to activate mechanosensitive channels in neurons, fibroblasts, and other titssues(32, 42, 43), its effects on LEC mechanosensitive channels remain unknown. Our RNA-sequencing of hLECs post-ultrasound results revealed upregulation of calcium-related pathways and multiple calcium signaling-related protein genes, suggesting that ultrasound may modulate calcium ion channels. The mechanosensitive ion channel Piezo1, capable of detecting forces below 10 pN, shows preferential calcium conductance in response to mechanical activation(29, 30). According to the Human Protein Atlas single-cell RNA database (as of October 2025), Piezo1 expression is most prominent in LECs compared to roughly 90 other human cell types. Previous studies have confirmed the distribution of Piezo1 on meningeal lymphatic endothelial cells, and genetic and pharmacological modulation of Piezo1 has demonstrated its importance in the development and maintenance of mLVs(11, 12). Therefore, we verified the response of the Piezo1 in LECs to ultrasound both in vivo and in vitro. Our in vitro live-cell calcium imaging experiments demonstrated that LIFU stimulation activated Piezo1 calcium channels in lymphatic endothelial cells, with channel activity persisting for approximately 2 min after stimulation ceased. While the high mechanosensitivity of Piezo1 underlies ultrasound’s ability to regulate calcium influx in mLECs, we also anticipate potential amplification effects through collaboration with other components. For instance, TRPV4 and SMPD3 may synergize with Piezo1 to potentiate ultrasound-mediated mechanoregulation in LECs(44, 45). However, the putative functional synergy between Piezo1 and other mechanosensitive components in lymphatic endothelial cells under ultrasound stimulation awaits experimental validation. Systematic investigation of these cooperative mechanotransduction pathways may elucidate their integrated regulatory mechanisms.

4-week LIFU stimulation enhanced mLVs-mediated drainage and expanded lymphatic coverage area. This sustained effect likely engages Piezo1-dependent pathways, given this channel’s established role as a critical regulator of lymphatic development and homeostasis(11, 12, 46, 47). To determine whether the chronic benefits of LIFU on mLVs are dependent on Piezo1 activation, we inhibited Piezo1 by intrathecal injection of the Piezo1 antagonist GsMTx4. Our results showed that inhibiting of Piezo1 abolished the beneficial effects of LIFU on mLVs drainage function, which is consistent with previous studies showing that genetic ablation or pharmacological inhibition of Piezo1 abolished the beneficial effects of Yoda1, a Piezo1 agonist, on mLVs(11, 12). This confirmed the importance of the Piezo1 ion channel for mLVs and also demonstrated that the benefits of LIFU are dependent on Piezo1 activation. It has been established that activation of Piezo1 increases lymphatic vessel permeability and stimulates pathways related to lymphatic development and maintenance, such as those involving VEGF-C and angiopoietin(12, 36). However, our study did not investigate the downstream signaling pathways mediated by Piezo1 following LIFU stimulation. Further exploration is warranted to fully elucidate the mechanisms underlying LIFU-mediated regulation of mLVs.

### Conclusion

We have developed an ultrasound stimulation protocol that precisely targets cranial meninges to non-invasively facilitate mLVs drainage. The target in situ peak negative pressure we used corresponds to a mechanical index (MI) of 0.5 (within the low-risk range of MI < 0.7) and is fully compliant with U.S. Food and Drug Administration safety guidelines(48), consistent with recent ultrasound neuromodulation studies(48, 49). Additionally, our stimulation protocol avoids direct excitation of the brain parenchyma and incorporates a relatively long ISI, thereby further improving its safety profile. CoS also presents distinct bony landmarks, which aids in reproducible clinical targeting. These safety attributes support the classification of this LIFU device as a nonsignificant risk device by local institutional review boards. If clinically validated, this LIFU strategy could offer a promising non-pharmacological and non-invasive adjunctive treatment for neurological disorders associated with impaired waste clearance in the central nervous system.

## Methods

### Animals

C57BL/6J strain mice were obtained from Jinzhihe Biotechnology Co., Ltd (Yangzhou, China). For the aging study, 18–19-month-old male wild-type mice were designated as the aged cohort, while 2–3-month-old males served as the young control group. T Alzheimer’s disease model was used: 9–10-month-old 5×FAD transgenic females. Corresponding control groups consisted of either non-transgenic littermates or sham-operated animals. Notably, amyloid plaque formation with concomitant glial activation becomes detectable in 5×FAD mice by 2 months of age, with cognitive deficits manifesting at approximately 5 months and neuronal degeneration in affected brain regions commencing around 6 months. All animals were maintained in standardized conditions featuring a 12-hour photoperiod, ambient temperature regulation (20–23°C), and relative humidity control (50–60%), with ad libitum access to rodent chow and water. Experimental procedures complied with International Council for Laboratory Animal Science (ICLAS) guidelines for laboratory animal welfare and were conducted under ethical approval (AMUWEC20242036) from the local institutional review board.

### In vivo mLVs low-intensity focused ultrasound (LIFU) stimulation protocol

Transcranial focused ultrasound (SY23WX013, Jie Lian, China), operating at a frequency of 500 kHz, was applied to dorsal mLVs (Figure 1A). We set output voltages of ultrasound to either 10V, 30V, 50V, or 70V. The measured values of spatial peak pulse average intensity (*I_SPPA_*) through the transcranial three-dimensional sound field measurement of ultrasound, after compensating for skull attenuation, were 0, 0.665, 1.578, 3.519, and 5.736W/cm^2^, respectively. Pulse repetition time (PRT) was set as 0.5ms. The duty cycle was set at 50%, with a 10-second inter-stimulus interval (ISI) for a total treatment duration of 20 minutes. Prior to ultrasound application, fur on the head was carefully removed using a chemical hair depilatory agent, to ensure optimal acoustic coupling. A collimator was then applied to the dorsal surface of the head, with ultrasound gel used to facilitate the transmission of ultrasonic waves. The mice were maintained under anesthesia with 1% isoflurane during the entire ultrasound treatment process. To prevent hypothermia, environmental heating was implemented to maintain body temperature within a normal physiological range.

### Transcranial three-dimensional sound field measurement of ultrasound

A three-dimensional scanning device was used to measure the entire sound field range. Computer software controlled the output voltage of the device. A conical collimator was installed on the transducer and the detached posterior fontanelle of a mouse skull was affixed onto the collimator using hot glue. The ultrasonic transducer (SY23WX013, focal length: 19.5 mm, frequency: 500 kHz) was driven to generate a sound field. A hydrophone (NDT, NM-0.5) received the ultrasonic signal and converted it into an electrical signal, which was displayed on an oscilloscope (DSO-X2024A). The motion controller (WNMC400) controlled the three-dimensional scanning device, enabling the hydrophone to move in the three-dimensional sound field, obtain the sound intensity at different positions, and thus measure the distribution of the sound field.

### Preparation of A**β**42 oligomers

Aβ42 oligomers were prepared as previously described(50). Briefly, 1 mg of Aβ42 peptide (Nanjing Peptide Biotech Ltd.) was dissolved in 0.2 mL of hexafluoroisopropanol (HFIP) and incubated at room temperature for 24 hours. After evaporating the HFIP under nitrogen gas, the peptide was re-dissolved in 100 μL of DMSO and then diluted with 1.6 mL of ultrapure water. The solution was incubated at 4°C for 24 hours and then centrifuged at 14,000 × g for 10 minutes at 4°C. The supernatant containing Aβ42 oligomers was collected.

### Hippocampus injections

We anesthetized mice with 1% isoflurane and immobilized their heads in a stereotaxic frame. After shaving and making a midline scalp incision, we administered 3 μL Aβ42 oligomers into each hippocampal CA1 region (coordinates from bregma: AP −2.3 mm, ML ±1.8 mm, DV −2.0 mm). The injection needle was maintained in position for 10 minutes before withdrawal to prevent solution backflow.

### Intra-cisterna magna (i.c.m.) injections

For Intra-cisterna magna injections, mice were anesthetized by intraperitoneal (i.p) injection of ketamine (80 mg/kg) and xylazine mix (20 mg/kg). The dorsal surface of the neck was shaved and cleaned. After making a 1.5 cm incision of the neck skin at midline, the muscle layers were retracted with forceps for the cisterna magna exposure. The desired solution was injected into the cisterna magna compartment (1 μL/min). For evaluation of lymphatic drainage, 5-10 μL of Alexa Fluor 647 conjugate ovalbumin (at 2 mg/mL, OVA-647, Thermo Fisher Scientific, USA) was administered using a Hamilton syringe. All i.c.m. injections were performed using a Hamilton syringe with a 33-G needle at a 135° angle. To prevent CSF leakage, the needle was maintained in position for an additional 10 minutes post-injection. Subsequently, the cervical skin and musculature were disinfected and surgically closed. The animals were then transferred to a temperature-regulated recovery surface until regaining consciousness, after which they were returned to their housing units.

### Intrathecal Injection (IT)

Intrathecal injections of 3μL GsMTx4(20 μM) were performed as described previously by Mestre et al(51). Mice were restrained in a prone position using a 50-mL conical tube for identification of the L5-L6 intervertebral space. A 30-gauge needle attached to a Hamilton syringe was advanced into the L4-L5 subarachnoid space. Successful intrathecal placement was confirmed by an immediate tail-flick reflex. Response times remained unchanged from pre-injection baselines following the procedure.

### IVIS imaging

Two hours post cisterna magna injection of OVA-647, euthanasia was performed via intraperitoneal administration of a lethal dose of Euthasol (10% v/v in saline). Following transcardial perfusion with cold saline, the dCLNs were excised. Fluorescence imaging was conducted using an IVIS Spectrum CT system (PerkinElmer), with subsequent quantification of bilateral dCLN signal intensity performed using Living Imaging software.

### Meningeal lymphatic vessels (mLVs) ablation

The Visudyne treatment was carried out following established methods(52). Briefly, mice were anesthetized using a combination of 1% isoflurane and dexmedetomidine. Subsequently, 10 μL of Visudyne solution (0.5 mg/mL, APExBIO, USA) was slowly administered via intracerebral injection (i.c.m.) at a controlled rate of 2.5 μL/min. Following a 15-minute incubation period, the drug was activated by exposure to a 689 nm cold laser (LSR689CP-3.6W-FC, LASEVER, China) applied to five predefined cranial locations: the injection site, the confluence of the sinuses, bilateral transverse sinuses, and the superior sagittal sinus. After the procedure, the scalp was surgically closed, and the animals were allowed to recover on a heating pad until fully awake before being returned to their housing.

### Laser speckle contrast imaging (LSCI)

After one month of treatment, mice were anesthetized with 1% isoflurane supplemented with dexmedetomidine. After surgical exposure of the skull through scalp and fascia resection, animals were secured in a head-fixation frame and positioned beneath an LSCI microscope (SIM BFI ZOOM, SIM Opto-Technology Co., Ltd) for cerebral blood flow assessment. A 785 nm excitation laser generated signals that were amplified and digitized for flow imaging. Continuous 3-minute recordings of relative blood flow (%) were obtained, with subsequent quantification of blood flow indices performed using manufacturer-provided software.

### Y-maze test

Spatial working memory was evaluated using a Y-maze (3 arms, 120° angles, 40 cm length) under dim light. Mice were habituated to the maze for 5 min on day 1. On day 2, mice were placed at the maze center and allowed 8 min of free exploration. Arm entries were recorded, with an “alternation” defined as sequentially entering all three arms without repetition. Object exploration was recorded using EthoVision XT software (version EV115, Noldus, Ltd.). Alternation Index= (Number of alternations) / (Total arm entries − 2) × 100%. Mice with <15 total arm entries were excluded. The maze was cleaned with 70% ethanol between trials. Spatial memory was evaluated using a Y-maze (3 arms, 120° angles, 40 cm length) with two arms blocked during habituation (day 1: 5 min). On day 2, mice explored the open arm for 5 min (training). After a 1-hour interval, all arms were opened, with one arm modified (e.g., texture change) as novel. Exploration time in each arm was recorded for 5 min. Object exploration was recorded using EthoVision XT software (version EV115, Noldus, Ltd.). Novel Arm Preference = (Novel Arm Time) / (Total Exploration Time) × 100%. Mice with <10 seconds of total exploration were excluded. The maze was cleaned with 70% ethanol between trials.

### Morris Water Maze (MWM) test

The MWM test was performed in a round pool (120 cm in diameter) containing opaque water at approximately 23°C, with a water depth of 30 cm. Various geometric shapes were positioned around the pool as visual cues. Over four consecutive training days, mice were placed in the water to locate a submerged platform (10 cm wide, positioned 1 cm beneath the water’s surface) within a 60-second time limit. If the mice did not find the platform within the allotted time, they were gently assisted onto it and allowed to remain there for 15 seconds. The latency to platform was recorded using EthoVision XT software (version EV115, Noldus, Ltd.) and the mean latency to platform of the four trials each day was calculated. During the probe trial on day 5, the platform was removed, and mice freely explored the pool for 60 seconds. Data were collected and analyzed, including quadrant dwell time (%), platform crossings, and latency.

### Human dermal lymphatic endothelial cell (LECs) culture

Lymphatic endothelial cells (LECs, cat.YIC-H161, ETHEPHON, Inc.) were grown in Endothelial Cell Medium (ECM, cat. #1001, ScienCell Research Laboratories, Inc.). The medium was supplemented with 5% fetal bovine serum (FBS), as well as vascular endothelial growth factor (VEGF), fibroblast growth factor (FGF), epidermal growth factor (EGF), and insulin-like growth factor (IGF). The cells were seeded onto wells and flasks that had been coated with Attachment Factor Protein (S006100, Gibco, Thermo Fisher Scientific). Subsequently, they were placed in a humidified incubator maintained at 37°C with a 5% CO[atmosphere for incubation.

### Cell siRNA transfection

Cells were seeded in 6-well plates at 70–80% confluency 24 hours prior to transfection. Piezo1 siRNA (20 nM) and Lipofectamine 3000 (mixed with siRNA at a volume-to-volume ratio of 1:1–1:2) were separately diluted in Opti-MEM. They were then combined and incubated at room temperature for 15–20 minutes to form complexes. The resulting mixture was added drop-by-drop to the cells in antibiotic-free medium. After 4–6 hours, the medium was replaced with complete medium. The efficiency of gene knockdown was assessed by qPCR 24–48 hours later.

### Calcium fluorescence imaging

To evaluate changes in intracellular calcium levels, cells were stained with the calcium-sensitive fluorescent probe Fluo4 AM (Beyotime, Shanghai, China; excitation wavelength: 494 nm, emission wavelength: 516 nm). First, cells cultured in 10-cm dishes were washed twice with Ca²□-free HBSS. Then, they were incubated with 4 μM Fluo-4AM for 20–30 minutes (the time was adjusted according to cell conditions) at 37 °C in a 5% CO□ environment. After the light-protected incubation, the dishes were rinsed three times with Ca²□-free HBSS at room temperature. Next, calcium oscillations were observed using an Olympus IX81 fluorescence microscope (Olympus Optical, Japan). Time-lapse images were taken at 5-second intervals. LIFU was applied with the following parameters: frequency of 0.5 MHz, *I_SPPA_*of 1.578 W/cm^2^, a duty cycle of 50%, and a stimulation time of 2 minutes.

### RNA extraction and library construction

Total RNA was extracted using TRIzol reagent (Invitrogen, Carlsbad, CA, USA) according to the standard procedure. The concentration and purity of the RNA were measured spectrophotometrically with a NanoDrop ND-1000 (NanoDrop, Wilmington, DE, USA), and its integrity was evaluated using an Agilent Bioanalyzer 2100 (Agilent, CA, USA). Only samples with an RNA Integrity Number (RIN) greater than 7.0 and distinct 18S/28S ribosomal bands on denaturing agarose gels were kept for further analysis. Poly(A)+ RNA was enriched from 1 μg of total RNA through two rounds of selection with Dynabeads Oligo(dT)25 beads (Thermo Fisher, USA). The purified mRNA was chemically fragmented using the Magnesium RNA Fragmentation Module (NEB #E6150) at 94°C for 5-7 minutes. First-strand cDNA synthesis was carried out using SuperScript II Reverse Transcriptase (Invitrogen #1896649). Subsequently, second-strand synthesis was performed with E. coli DNA polymerase I (NEB #M0209), RNase H (NEB #M0297), and dUTP (Thermo Fisher #R0133) to produce U-labeled double-stranded DNA (dsDNA). The blunt-ended cDNA fragments were A-tailed and then ligated to indexed adapters with T-overhangs. Size selection (300 ± 50 bp) was done using AMPureXP beads. Uracil residues were removed by UDG (NEB #M0280) treatment before PCR amplification. The PCR conditions were as follows: 95°C for 3 minutes; 8 cycles of 98°C for 15 seconds, 60°C for 15 seconds, and 72°C for 30 seconds; and a final extension at 72°C for 5 minutes. The final libraries were quantified and pooled for 2×150 bp paired-end sequencing on an Illumina Novaseq 6000 (LC − Bio Technology, Hangzhou, China) following the manufacturer’s instructions.

### Bioinformatics analysis of RNA-seq

We employed fastp (v0.23.2, OpenGene) to process raw sequencing reads. Using its default parameters, adapter sequences, low-quality bases (Q < 20), and ambiguous nucleotides were removed. After filtering, the same tool was used to re-evaluate the read quality metrics.For read mapping, we utilized Ensembl_v112 (ftp://ftp.ensembl.org/pub/release-112/fasta/homo_sapiens/dna/) to map reads to the Homo sapiens GRCh38 reference genome and Ensembl_v107 (ftp://ftp.ensembl.org/pub/release-107/fasta/mus_musculus_c57bl6nj/dna/) for mapping reads to the mus musculus c57bl6nj reference genome. Next, we assembled the mapped reads of each sample using StringTie (https://ccb.jhu.edu/software/stringtie) with default settings. Subsequently, gffcompare (https://github.com/gpertea/gffcompare/) was applied to merge all transcriptomes from all samples and reconstruct a comprehensive transcriptome. Once the final transcriptome was generated, StringTie was used to estimate the expression levels of all transcripts. Specifically, for mRNAs, StringTie calculated the FPKM values (FPKM = [total_exon_fragments / mapped_reads(millions) × exon_length(kB)]). To identify differentially expressed mRNAs, we used the R package edgeR (https://bioconductor.org/packages/release/bioc/html/edgeR.html). MRNAs with a fold-change greater than 2 or less than 0.5 and a p-value less than 0.05 from a parametric F-test comparing nested linear models were selected.

### Tissue collection and processing

Mice were euthanized via intraperitoneal injection of a lethal dose of Euthasol (10% v/v in physiological saline), followed by transcardial perfusion with ice-cold phosphate-buffered saline (PBS, pH 7.4) containing heparin (10 U/mL). After removing the skin and muscles, the intact head was excised and fixed in 4% paraformaldehyde (PFA) at 4°C for 24 hours. The mandible and nasal bones were carefully removed, and the skull cap was excised using fine curved surgical scissors (Fine Science Tools). The cut was made in a clockwise direction, starting and ending near the hook posterior to the right tympanic membrane. The dissected skull cap was then stored in PBS containing 0.02% sodium azide at 4°C. For meningeal extraction, the dura mater and arachnoid mater were gently separated from the remaining skull using Dumont #5 and #7 forceps (Fine Science Tools) and preserved in azide-supplemented PBS at 4°C. Alternatively, the skull was bisected along the sagittal midline, and after brain removal, the meningeal-covered skull fragments were similarly stored. The brain, dCLNs, and dorsal cranial bone (with attached dura) underwent additional fixation in 4% PFA for 24 hours (48 hours total). After PBS washing, the brain was cryoprotected in 30% sucrose, embedded in Tissue-Plus® O.C.T. compound, and sectioned at 30 μm thickness using a Leica cryostat. Sections were stored in azide-PBS at 4°C until further processing.

For immunofluorescence staining, brain sections, CLN sections, and meninges were initially rinsed with PBS. They were then blocked for 2 hours at room temperature (RT) in a PBS solution containing 0.3% Triton X-100 (Sigma, USA) and 5% bovine serum albumin (BSA). After the blocking step, the tissues were incubated overnight at 4 °C with the following primary antibodies at the indicated dilutions: rabbit anti-LYVE-1 (Cell Signaling, 67538S, 1:400), goat anti-CD31 (R&D, AF3628, 1:200), goat anti-Iba1 (Wako, 011-27991, 1:800), mouse anti-NeuN (Cell Signaling, 94403S, 1:400), rabbit anti-amyloid-β37–42 (Cell Signaling, clone D54D2, 1:300), rabbit anti-GFAP (Cell Signaling, 12389S, 1:400), rabbit anti-HSP 70 (Proteintech, 10995-1-AP, 1:400), and rabbit anti-c-Fos (Cell Signaling, 2250s, 1:400). Next, the tissues were washed three times with PBS. Subsequently, they were incubated for 2 hours in the dark at RT with Alexa Fluor 488/555/647/594 donkey anti-goat/rabbit/mouse IgG (H + L) secondary antibodies in a PBS solution with 0.3% Triton X-100. The specific secondary antibodies employed were: Alexa Fluor 555 donkey anti-goat IgG (H + L) secondary antibody (Abcam, ab150129, 1:500), Alexa Fluor 488 rabbit anti-goat IgG (H + L) secondary antibody (Abcam, ab150073, 1:500), and Alexa Fluor 594-conjugated affinipure donkey anti-mouse IgG (H+L) (Jackson, 715-585-150, 1:800). After incubation with secondary antibodies, the tissues were washed three additional times and then incubated with DAPI for 5 minutes at RT. Finally, images were taken using a confocal microscope (Ziss 880). All measurements were performed by an experimenter unaware of the sample identities. Microsoft Excel was used to calculate the average values for each experiment, and statistical analysis was performed using GraphPad Prism 8.3.4.

### Statistical analysis

To achieve randomization, animals from various cages were distributed evenly into the same experimental groups. Throughout data collection and analysis, researchers were unaware of the group assignments for the mice. Sample sizes were chosen based on previous experiments(7, 37). Statistical methods were not reapplied to predetermine sample sizes. Normal distribution of data was first checked with the Anderson-Darling, D’Agostino, and Shapiro–Wilk normality tests. For statistical evaluation, two-group comparisons were performed using two-tailed unpaired Student’s t tests, while multiple-group comparisons were assessed with one-way followed by S Bonferroni’s post hoc test or HolmSidak’s post hoc test. Longitudinal data (day × treatment) were analyzed using repeated-measures two-way ANOVA with Bonferroni post hoc testing. A significance threshold of *P* < 0.05 was adopted. Results are expressed as mean ± standard error of the mean (SEM). All statistical analyses were carried out using GraphPad Prism (v8.3.4, GraphPad Software, San Diego, CA).

## Acknowledgements

We are grateful to the Hou lab members for their insightful discussions and technical support throughout this study.

## Author contributions

X. Xu, F. Zhou, X. Li and J. Hou designed the study; X. Xu, T. Jiang, C. Zhu, Y. Shu and Y. Tang performed experiments; X. Xu, T. jiang and C. Zhu contributed to data analysis; X. Xu analyzed data and wrote the manuscript; F. Zhou, X. Li and J. Hou revised the manuscript, X. Li and J. Hou oversaw the project. All the authors read and approved the manuscript.

## Availability of data and materials

The data that support the findings of this study are available from the corresponding author upon reasonable request. BioProject and associated SRA metadata are available at https://dataview.ncbi.nlm.nih.gov/object/PRJNA1259799?reviewer=h8t81c2e99cv5si 9287vujp9a1 in read-only format.

## Declarations

### Ethics approval and consent to participate

Mouse experiments were approved by Laboratory Animal Welfare and Ethics Committee of the Army Medical University (AMUWEC20242036).

### Consent for publication

No applicable

## References

1. 2024 Alzheimer’s disease facts and figures. Alzheimer’s & Dementia 20, 3708–3821 (2024).

2. P. Scheltens, et al., Alzheimer’s disease. Lancet 397, 1577–1590 (2021).

3. J. Kipnis, Multifaceted interactions between adaptive immunity and the central nervous system. Science 353, 766–771 (2016).

4. A. Louveau, T. H. Harris, J. Kipnis, Revisiting the Mechanisms of CNS Immune Privilege. Trends Immunol 36, 569–577 (2015).

5. A. Aspelund, et al., A dural lymphatic vascular system that drains brain interstitial fluid and macromolecules. J Exp Med 212, 991–999 (2015).

6. A. Louveau, et al., Structural and functional features of central nervous system lymphatic vessels. Nature 523, 337–341 (2015).

7. S. Da Mesquita, et al., Functional aspects of meningeal lymphatics in ageing and Alzheimer’s disease. Nature 560, 185–191 (2018).

8. S. Da Mesquita, et al., Meningeal lymphatics affect microglia responses and anti-Aβ immunotherapy. Nature 593, 255–260 (2021).

9. D. Choi, et al., Piezo1-Regulated Mechanotransduction Controls Flow-Activated Lymphatic Expansion. Circ Res 131, e2–e21 (2022).

10. L. Planas-Paz, et al., Mechanoinduction of lymph vessel expansion. EMBO J 31, 788–804 (2012).

11. M. J. Matrongolo, et al., Piezo1 agonist restores meningeal lymphatic vessels, drainage, and brain-CSF perfusion in craniosynostosis and aged mice. J Clin Invest 134, e171468 (2023).

12. D. Choi, et al., Piezo1 regulates meningeal lymphatic vessel drainage and alleviates excessive CSF accumulation. Nat Neurosci 27, 913–926 (2024).

13. J. Du, et al., The mechanosensory channel PIEZO1 functions upstream of angiopoietin/TIE/FOXO1 signaling in lymphatic development. J Clin Invest 134, e176577 (2024).

14. K. Nonomura, et al., Mechanically activated ion channel PIEZO1 is required for lymphatic valve formation. Proc Natl Acad Sci U S A 115, 12817–12822 (2018).

15. N. Lipsman, K. Hynynen, R. Chen, A. M. Lozano, Transcranial focused ultrasound in the human brain. Neuron 114, 601–621 (2026).

16. V. Krishna, F. Sammartino, A. Rezai, A Review of the Current Therapies, Challenges, and Future Directions of Transcranial Focused Ultrasound Technology: Advances in Diagnosis and Treatment. JAMA Neurol 75, 246–254 (2018).

17. K. Lee, T. Y. Park, W. Lee, H. Kim, A review of functional neuromodulation in humans using low-intensity transcranial focused ultrasound. Biomed Eng Lett 14, 407–438 (2024).

18. R. Beisteiner, M. Hallett, A. M. Lozano, Ultrasound Neuromodulation as a New Brain Therapy. Adv Sci (Weinh) 10, e2205634 (2023).

19. C.-H. Wu, et al., Very Low-Intensity Ultrasound Facilitates Glymphatic Influx and Clearance via Modulation of the TRPV4-AQP4 Pathway. Adv Sci (Weinh) 11, e2401039 (2024).

20. S. Choi, et al., Transcranial focused ultrasound stimulation enhances cerebrospinal fluid movement: Real-time in vivo two-photon and widefield imaging evidence. Brain Stimul 17, 1119–1130 (2024).

21. S.-S. Yoo, et al., Non-invasive enhancement of intracortical solute clearance using transcranial focused ultrasound. Sci Rep 13, 12339 (2023).

22. M. Aryal, et al., Noninvasive ultrasonic induction of cerebrospinal fluid flow enhances intrathecal drug delivery. J Control Release 349, 434–442 (2022).

23. Y. Lee, et al., Improvement of glymphatic–lymphatic drainage of beta-amyloid by focused ultrasound in Alzheimer’s disease model. Sci Rep 10, 16144 (2020).

24. M. M. Azadian, et al., Clearance of intracranial debris by ultrasound reduces inflammation and improves outcomes in hemorrhagic stroke models. Nat Biotechnol (2025). 10.1038/s41587-025-02866-8.

25. C. Pasquinelli, L. G. Hanson, H. R. Siebner, H. J. Lee, A. Thielscher, Safety of transcranial focused ultrasound stimulation: A systematic review of the state of knowledge from both human and animal studies. Brain Stimul 12, 1367–1380 (2019).

26. L. C. D. Smyth, et al., Identification of direct connections between the dura and the brain. Nature 627, 165–173 (2024).

27. J. H. Ahn, et al., Meningeal lymphatic vessels at the skull base drain cerebrospinal fluid. Nature 572, 62–66 (2019).

28. P. P. Qin, et al., The effectiveness and safety of low-intensity transcranial ultrasound stimulation: A systematic review of human and animal studies. Neurosci Biobehav Rev 156, 105501 (2024).

29. B. Coste, et al., Piezo1 and Piezo2 are essential components of distinct mechanically activated cation channels. Science 330, 55–60 (2010).

30. J. Wu, R. Goyal, J. Grandl, Localized force application reveals mechanically sensitive domains of Piezo1. Nat Commun 7, 12939 (2016).

31. J. Zhu, et al., The mechanosensitive ion channel Piezo1 contributes to ultrasound neuromodulation. Proc Natl Acad Sci U S A 120, e2300291120 (2023).

32. Z. Jiang, et al., Low-Frequency Ultrasound Sensitive Piezo1 Channels Regulate Keloid-Related Characteristics of Fibroblasts. Adv Sci (Weinh) 11, e2305489 (2024).

33. F. Zheng, et al., Low-intensity pulsed ultrasound promotes the osteogenesis of mechanical force-treated periodontal ligament cells via Piezo1. Front Bioeng Biotechnol 12, 1347406 (2024).

34. S. Sun, et al., Efficient generation of human NOTCH ligand-expressing haemogenic endothelial cells as infrastructure for in vitro haematopoiesis and lymphopoiesis. Nat Commun 15, 7698 (2024).

35. N. Morooka, et al., Angpt1 binding to Tie1 regulates the signaling required for lymphatic vessel development in zebrafish. Development 151, dev202269 (2024).

36. J. Du, et al., The mechanosensory channel PIEZO1 functions upstream of angiopoietin/TIE/FOXO1 signaling in lymphatic development. J Clin Invest 134, e176577 (2024).

37. M. Wang, et al., Non-invasive modulation of meningeal lymphatics ameliorates ageing and Alzheimer’s disease-associated pathology and cognition in mice. Nat Commun 15, 1453 (2024).

38. S. Da Mesquita, et al., Meningeal lymphatics affect microglia responses and anti-Aβ immunotherapy. Nature 593, 255–260 (2021).

39. J.-H. Yoon, et al., Nasopharyngeal lymphatic plexus is a hub for cerebrospinal fluid drainage. Nature 625, 768–777 (2024).

40. J. H. Ahn, et al., Meningeal lymphatic vessels at the skull base drain cerebrospinal fluid. Nature 572, 62–66 (2019).

41. L. C. D. Smyth, et al., Identification of direct connections between the dura and the brain. Nature 627, 165–173 (2024).

42. Z. Qiu, et al., The Mechanosensitive Ion Channel Piezo1 Significantly Mediates In Vitro Ultrasonic Stimulation of Neurons. iScience 21, 448–457 (2019).

43. S. Yoo, D. R. Mittelstein, R. C. Hurt, J. Lacroix, M. G. Shapiro, Focused ultrasound excites cortical neurons via mechanosensitive calcium accumulation and ion channel amplification. Nat Commun 13, 493 (2022).

44. J. Shi, et al., Sphingomyelinase Disables Inactivation in Endogenous PIEZO1 Channels. Cell Rep 33, 108225 (2020).

45. S. M. Swain, R. A. Liddle, Piezo1 acts upstream of TRPV4 to induce pathological changes in endothelial cells due to shear stress. J Biol Chem 296, 100171 (2021).

46. K. Nonomura, et al., Mechanically activated ion channel PIEZO1 is required for lymphatic valve formation. Proc Natl Acad Sci U S A 115, 12817–12822 (2018).

47. D. Choi, et al., Piezo1 incorporates mechanical force signals into the genetic program that governs lymphatic valve development and maintenance. JCI Insight 4, e125068, 125068 (2019).

48. E. Martin, et al., ITRUSST consensus on standardised reporting for transcranial ultrasound stimulation. Brain Stimul 17, 607–615 (2024).

49. D. Attali, et al., Three-layer model with absorption for conservative estimation of the maximum acoustic transmission coefficient through the human skull for transcranial ultrasound stimulation. Brain Stimul 16, 48–55 (2023).

50. Y. Wu, et al., Borneol-driven meningeal lymphatic drainage clears amyloid-β peptide to attenuate Alzheimer-like phenotype in mice. Theranostics 13, 106–124 (2023).

51. C. Mestre, T. Pélissier, J. Fialip, G. Wilcox, A. Eschalier, A method to perform direct transcutaneous intrathecal injection in rats. J Pharmacol Toxicol Methods 32, 197–200 (1994).

52. M. Wang, et al., Non-invasive modulation of meningeal lymphatics ameliorates ageing and Alzheimer’s disease-associated pathology and cognition in mice. Nat Commun 15, 1453 (2024).

